# Structural basis of epitope recognition by anti-alpha synuclein antibodies MJFR14-6-4-2

**DOI:** 10.1101/2023.10.27.564328

**Authors:** Ilva Liekniņa, Teodors Panteļejevs, Alons Lends, Lasse Reimer, Kristaps Jaudzems, Aadil El-Turabi, Hjalte Gram, Poul Henning Jensen, Kaspars Tārs

## Abstract

Intraneuronal α-synuclein inclusions in the brain are hallmarks of so-called Lewy body diseases - Parkinson’s disease and Dementia with Lewy bodies. Lewy bodies are cytoplasmic inclusions, containing mainly aggregated α-synuclein together with some other proteins including ubiquitin, neurofilament protein, and alpha B crystallin. In its monomeric form, α-synuclein is predominantly localized in nerve terminals, regulating neuronal transmission and synaptic vesicle trafficking. Monomeric α-synuclein lacks a well-defined three-dimensional structure and is considered an intrinsically disordered protein. However, in diseased cells α-synuclein aggregates into oligomeric and fibrillar amyloid species, which can be detected using aggregate-specific antibodies. Here we investigate the aggregate specificity of rabbit monoclonal MJFR14-6-4-2 antibodies, preferentially recognizing aggregated α-synuclein species. We conclude that partial masking of epitope in unstructured monomer in combination with a high local concentration of epitopes instead of distinct epitope conformation is the main reason for apparent selectivity towards various aggregates, including oligomers, fibrils, and artificial virus-like particle constructs bearing multiple copies of the MJFR14-6-4-2 epitope. Based on the structural insight, we were able to express mutant α-synuclein that when fibrillated are unable to bind MJFR14-6-4-2. Using these “stealth” fibrils as a tool for seeding cellular α-synuclein aggregation, provides superior signal/noise ratio for detection of cellular α-synuclein aggregates by MJFR14-6-4-2 immunocytochemistry. Our data provide a molecular level understanding of specific recognition of toxic amyloid oligomers, which is critical for the development of inhibitors against synucleinopathies.

## Introduction

Alpha-synuclein (α-syn) is a protein that is mainly expressed in presynaptic terminals of neurons and plays a role in synaptic vesicle trafficking and neurotransmission [1–3]. However, α-syn is also implicated in the pathogenesis of Parkinson’s disease (PD) and other synucleinopathies, such as dementia with Lewy bodies, multiple system atrophy, pure autonomic failure, REM sleep behavior disorder [4]. These diseases are characterized by the accumulation and spreading of abnormal aggregates of α-syn, known as Lewy bodies and Lewy neurites in neurons, or glial cytoplasmic inclusions in glial cells [5, 6]. The aggregation of α-syn is influenced by various factors, such as mutations, post-translational modifications, oxidative stress, and interactions with other proteins or lipids [7, 8]. The aggregated forms of α-syn are thought to impair various cellular functions and pathways, such as mitochondrial dynamics, lysosomal degradation, calcium homeostasis, and inflammation, leading to neuronal dysfunction and death [4, 9]. The deposition of α-syn also correlates with synucleinopathies’ clinical features and progressions, such as cognitive impairment, motor symptoms, autonomic dysfunction, and sleep disturbances [10, 11]. Therefore, α-syn is a major therapeutic target for treating and preventing synucleinopathies. Several strategies have been developed or are under investigation to modulate α-syn levels, aggregation, toxicity, or clearance. These include immunotherapy, gene therapy, small molecule inhibitors or modulators of α-syn aggregation or degradation, and neuroprotective agents [12–14]. However, there are still many challenges and limitations to overcome before these therapies can be translated into clinical practice. These include the lack of reliable biomarkers for early diagnosis and disease monitoring, the heterogeneity and complexity of synucleinopathies, the potential side effects or adverse reactions of the therapies, and the difficulty of delivering the therapies across the blood-brain barrier [4, 15, 16].

The rabbit monoclonal MJFR14-6-4-2 is suggested to be a conformation-specific antibody that recognizes and binds aggregated forms of α-synuclein. These antibodies were generated in a collaboration between the Dandrite & Department of Biomedicine, University of Aarhus, Denmark, Epitomics Inc., and The Michael J. Fox Foundation, and marketed by Abcam [17]. One application of the MJFR14-6-4-2 antibody is to measure α-syn oligomers, which are small and soluble aggregates of α-syn that are considered to be the most toxic species in synucleinopathies. MJFR14-6-4-2 antibody has been used to perform an enzyme-linked immunosorbent assay for quantification of α-syn oligomers in cell and animal models of PD [17]. This demonstrates its potential to evaluate the effects of different factors or treatments on α-syn oligomerization, such as mutations, post-translational modifications, oxidative stress, inflammation, and pharmacological agents, as well as to detect α-syn aggregates in various biological fluids or extracts, such as cerebrospinal fluid, plasma, saliva, urine, or brain homogenates. This technique can be used to compare the levels of α-syn aggregates between different groups or treatments, such as healthy controls versus patients or animals with synucleinopathies [17, 18]. Another application of MJFR14-6-4-2 antibody is to visualize α-syn aggregates in human and animal tissue sections or cell cultures using immunohistochemistry or immunocytochemistry techniques [19, 20]. Finally, the antibody can be used in the diagnosis of PD biomarkers in human blood samples [21]. However, the specificity of MJFR14-6-4-2 and other anti-α-synuclein antibodies towards oligomeric and fibrillar forms of protein has been questioned previously [22], necessitating further research in the area.

Virus-like particles (VLPs) are self-assembled structures that mimic the morphology and antigenicity of natural viruses but lack the viral genome and are therefore non-infectious and safe. VLPs can be used as platforms for displaying foreign epitopes or antigens and thus serve as attractive tools for vaccine development and diagnostics [23]. Antigens presented on VLPs are so immunogenic that antibody responses can be elicited against self-antigens, thus overcoming the tolerance mechanisms in the context of these recombinant constructs [24]. Multiple repetitive copies of epitopes in VLPs induce cross-linking of B-cell receptors, greatly enhancing B-cell response. Therefore, recombinant VLP-based vaccines could replace therapeutic monoclonal antibodies as treatments for some diseases [25]. Significantly, VLPs can be used as a scaffold to multimerize epitopes of given antigens in a more well-ordered manner. This property allows the use of VLPs to investigate the effects of epitope multimerization in the absence of other parts of the antigen. A feature we propose can be exploited when studying certain aggregate species.

Among the various types of VLPs, those derived from ssRNA phages have several advantages, such as high stability, easy production and purification, low cost, and versatility [26]. The virus-like particles formed by the AP205 phage coat protein are unique in that both the C-and N-termini are exposed on the surface, therefore any C-or N-terminal fused antigens are accessible to the immune system [27].

In this study, we explore epitope recognition of MJFR14-6-4-2 antibodies using protein x-ray crystallography, NMR, and binding studies using our VLP library with exposed α-syn and develop a novel preformed alpha-synuclein fibrils extended by a C-terminal glycine not detectable by MJFR14-6-4-2 to study the cell biology of seeded cellular alpha-synuclein aggregation by MJFR14-6-4-2-based microscopy without interference from the exogenous alpha-synuclein fibrils.

## Results and Discussion

### Mapping of the MJFR14-6-4-2 epitope

Although it was known from previous experiments that the structural basis for the MJFR14-6-4-2 epitope lies in the C-terminal part of α-synuclein, the exact epitope was uncertain. Therefore, for mapping this and other α-synuclein epitopes we created a library of partially overlapping 14-mer peptides (AP-1 to AP-19), covering the entire α-synuclein sequence and displaying on bacteriophage AP205 virus-like particles (VLPs) as C-terminal fusion proteins with AP205 coat protein as shown in Figure 1a. In the library, we also included 10 residue-long C-terminal peptide PD3, immunization of which has previously led to the formation of aggregate-specific α-synuclein antibodies [28]. As expected, in direct ELISA MJFR14-6-4-2 reacted strongly with VLPs bearing the C-terminal α-synuclein sequence (AP-19, see Figure 1b). Subsequently, we constructed VLPs with progressively shorter C-terminal α-synuclein sequences (AP-19a to AP-19i). The results revealed that the C-terminal sequence EPEA corresponding to residues 137-140 constitutes the minimal epitope (Figure 1c). Furthermore, the VLP construct bearing an additional glycine in the C-terminus (AP-19k, epitope sequence EPEAG) was not recognized by MJFR14-6-4-2 antibodies, suggesting that the free carboxyl group of terminal alanine in α-synuclein is essential for antibody recognition.

**Figure 1.**
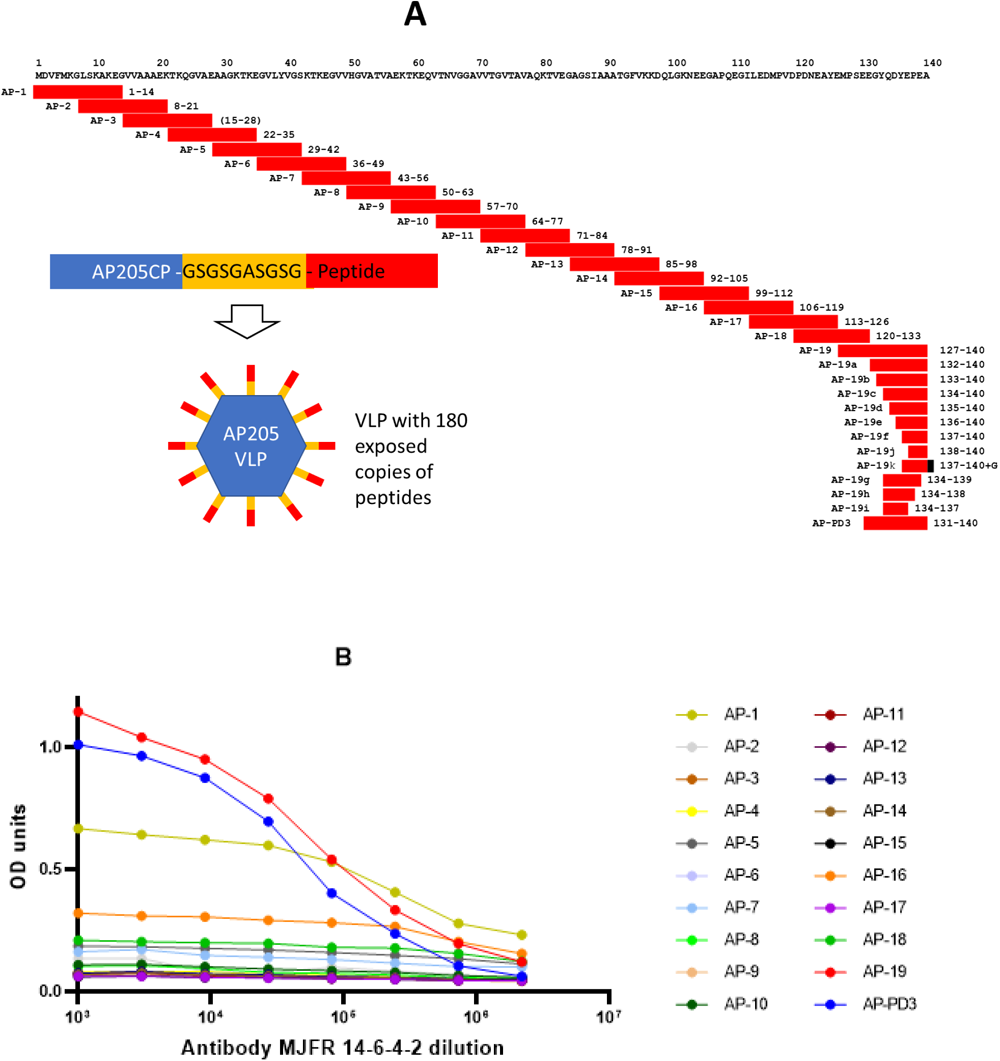
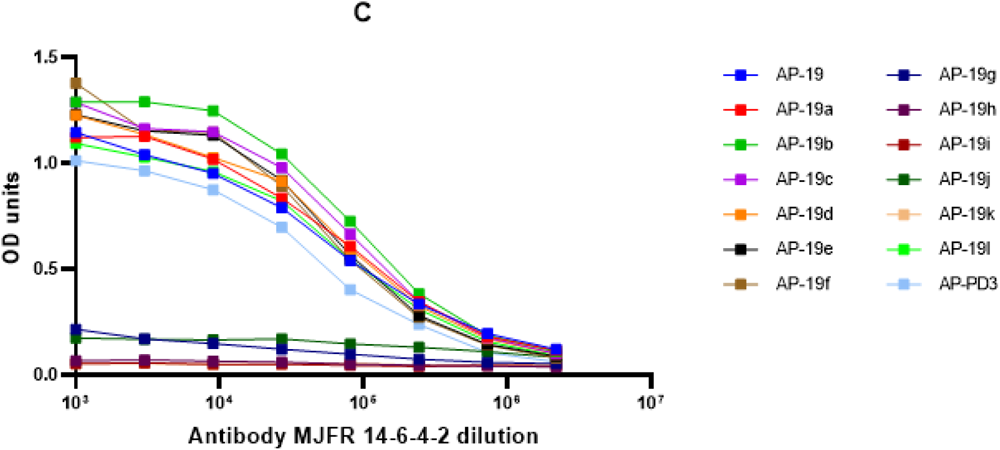
Mapping of epitope by ELISA. (A) Alpha-synuclein-AP205 VLP constructs for epitope mapping. For initial screening, 19 partially overlapping peptides along with glycine-serine linker were fused to the C-terminus of bacteriophage AP205 coat protein, resulting in recombinant VLPs. For fine mapping of the C-terminal epitope, further constructs 19a-j were produced as indicated. PD3 sequence was used in the construction of vaccine candidates previously [28].; (B) initial and (C) fine mapping of epitope

During the initial mapping, we observed that the N-terminal α-synuclein peptide (AP-1) displayed on VLPs also bound to MJFR14-6-4-2 antibodies, although with lower capacity as judged from the approximate 2-fold lower OD values observed in the ELISA compared to C-terminal peptide. However, further competition ELISA experiments revealed that AP-1 is unable to out-compete EPEA-epitope-containing constructs, therefore the N-terminal sequence of α-synuclein is not likely to comprise part of the epitope.

### Binding of antibodies to an epitope in different monomeric and multimeric species

We conducted ELISA competitions to compare epitope recognition within VLPs, α-synuclein monomers, and soluble oligomers formed spontaneously when lyophilized α-synuclein is reconstituted and purified by gel filtration. Oligomers were absorbed on plates and incubated with antibodies pre-treated with different competitors: peptide, α-synuclein species, or VLPs with EPEA epitope. α-Synuclein monomers showed no competition (Figure 2a), consistent with prior findings. Oligomers in solution outcompeted plate-absorbed oligomers. VLPs with EPEA epitopes exhibited the most effective competition. Plate-absorbed VLPs were mainly outcompeted by the same VLPs in solution; oligomers showed limited competition (Figure 2b), and monomers showed none. While initial ELISA suggested comparable binding of VLPs with minimal versus longer EPEA epitopes, competition studies favoured slightly longer exposed epitopes (Figure 3c).

**Figure 2.**
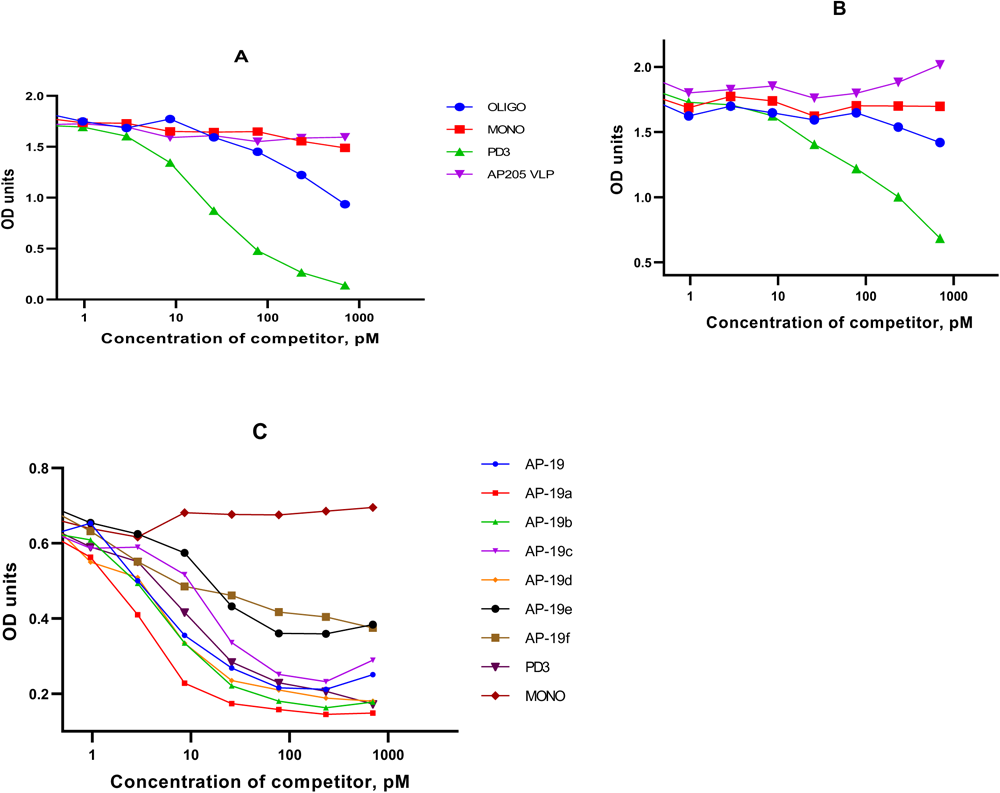
Competition ELISAs with antibodies. (A) ELISA with MJFR 14-6-4-2 antibodies, oligomers adsorbed on plates; (B) ELISA with MJFR 14-6-4-2 antibodies, PD3 VLPs adsorbed on plates; (C) same as (B) but comparison of competitivity of VLPs with epitopes of various lengths is shown.

**Figure 3.**
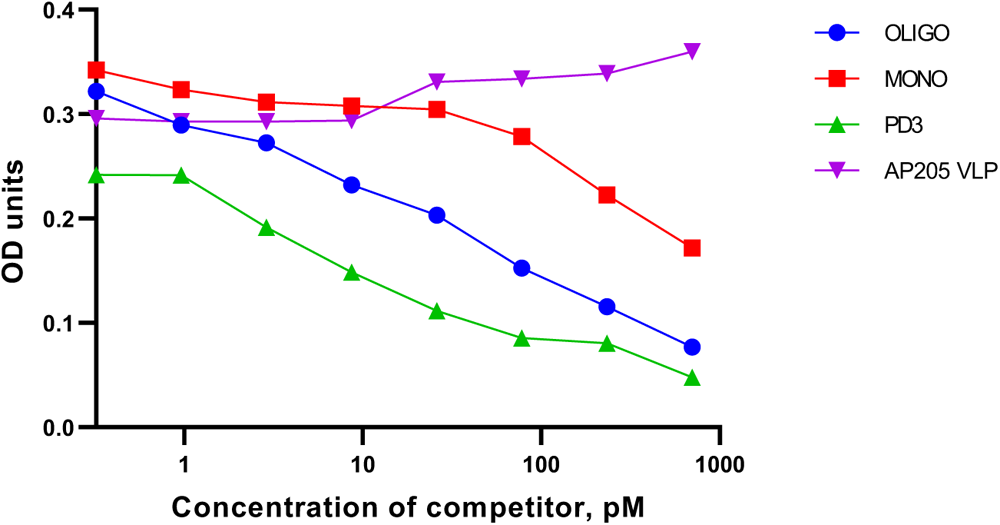
Competition ELISA with Fab fragments.

To see if the preferential binding to multimeric species is from an increased avidity effect due to the dual valency nature of IgG, we repeated competition experiments with prepared Fab fragment of MJFR14-6-4-2 antibody. As can be seen from Fig 3, monomeric α-synuclein species were able to some extent to compete with fibrils. However, oligomers and VLPs were still significantly better competitors. This demonstrates that increased reactivity with aggregated species is not merely due to the dimeric nature of IgG, but a more optimal presentation of the C-terminal tetrapeptide sequence.

The C-terminal part of α-synuclein has been demonstrated to engage in long-range interactions with regions in the middle of the protein. We hypothesised that such long range interactions are hiding the C-terminal MJFR14-6-4-2 epitope when α-synuclein is in its monomeric state. To test this hypothesis we wanted to explore the impact of the alpha helically structured that is induced in the N-terminal 95 amino acid residues upon binding to negatively charged liposomes [29, 30]. This change could prevent the long range interactions and thus expose the MJFR14-6-4-2 epitope in the liposome bound monomer.

By treating monomeric α-synuclein with liposomes, we indeed observed that at high α-synuclein concentrations, there was increased competition with VLP-exposed epitope compared to α-synuclein not treated with liposomes (Figure 4a). We confirmed by CD spectroscopy that our liposome-treated α-synuclein adopts alpha-helical conformation (Figure 4b). This suggests that the C-terminal epitope is somewhat unmasked in the alpha-helical conformation of α-synuclein. Alternatively, liposomes with embedded α-synuclein monomers and exposed C-termini may to some extent mimic VLPs with exposed C-terminal epitopes.

**Figure 4.**
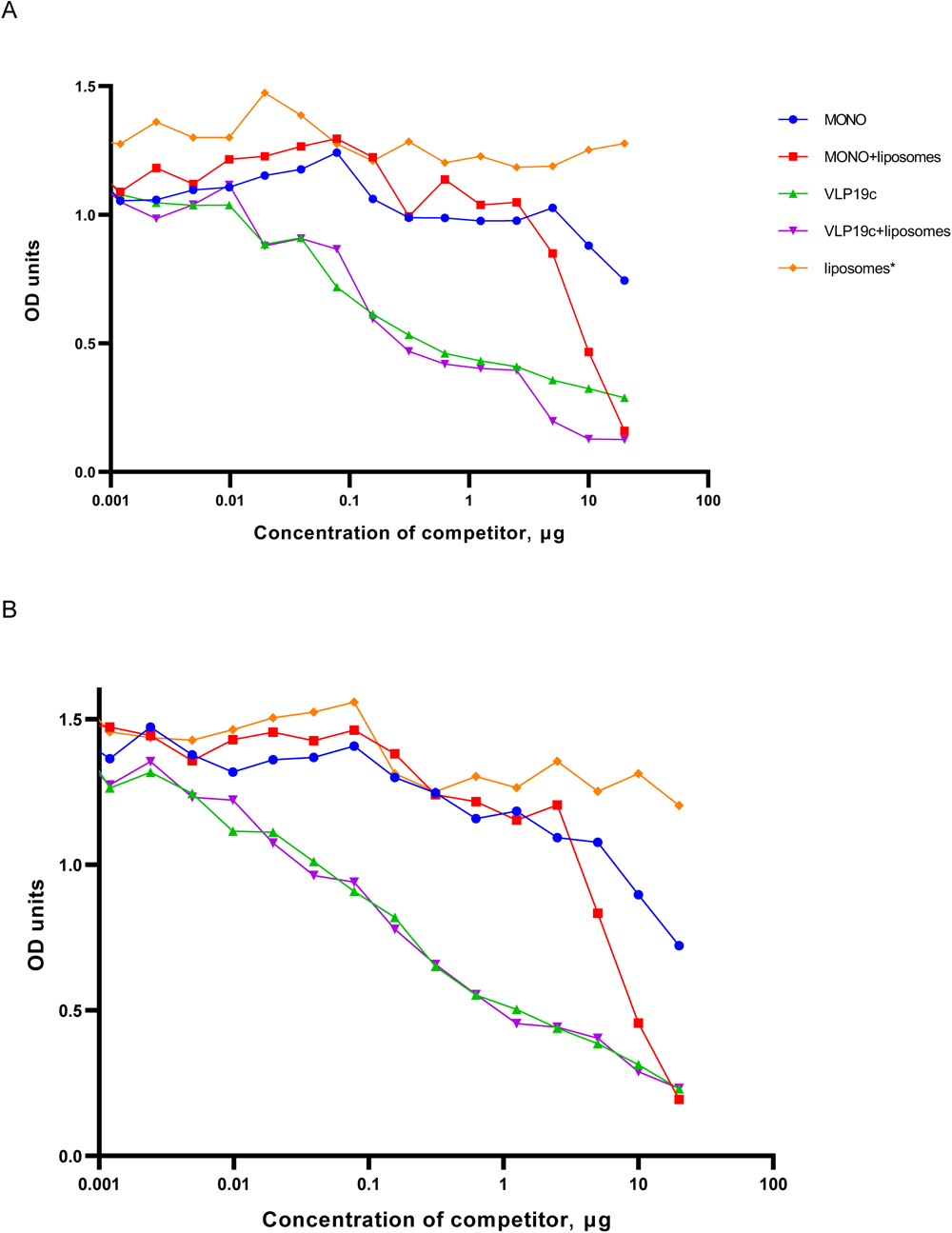
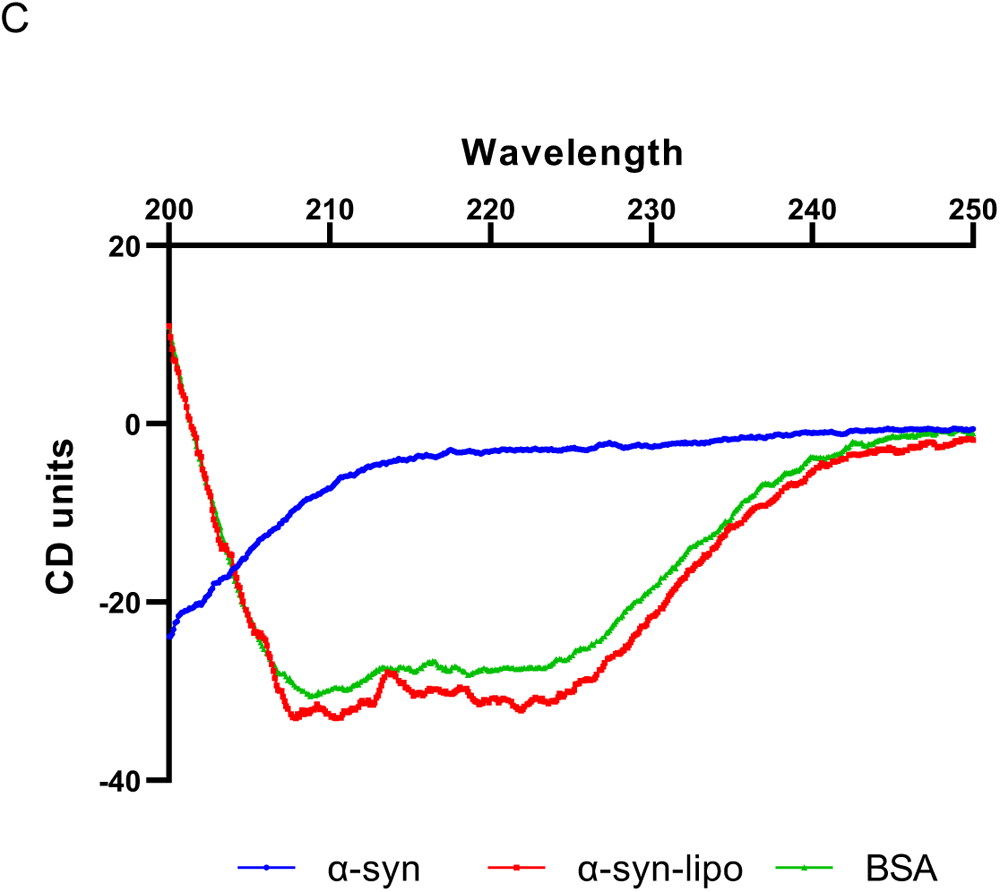
Effects of liposomes on the structure of α-synuclein and its binding to antibodies. (A) Competition ELISA with α-synuclein adsorbed on plates (B) Competition ELISA with VLP19c adsorbed on ELISA plates (C) CD curves of α-synuclein before and after the addition of liposomes. The curve of α-helical bovine serum albumin (BSA) is shown for comparison.

The enhanced reactivity of MJFR14-6-4-2 antibodies and their Fab fragments towards multimeric species was further confirmed by grating-coupled interferometry (GCI). We employed the waveRAPID® method [31] to measure binding kinetics. MJFR14-6-4-2 antibody was immobilised on a protein-A/G sensor chip, and α-synuclein species were used as analytes (Figure 5A-C). As anticipated, the α-synuclein monomers exhibited the lowest affinity towards the antibody (K_D_ = 0.530 μM) with fast dissociation. Whereas both oligomers and EPEA-displaying VLP did not dissociate from the surface, suggesting much stronger interactions and rendering the fitting of a kinetic model impossible. These observations are in line with a previous report describing the binding kinetics of MJFR14-6-4-2 and other antibodies to α-synuclein monomers and oligomers [22]. There, authors had similarly immobilised the antibody on an SPR chip surface and obtained a low-micromolar KD value for monomer binding with rapid dissociation, while demonstrating a much higher affinity for oligomers, with no discernable dissociation curvature.

**Figure 5.**
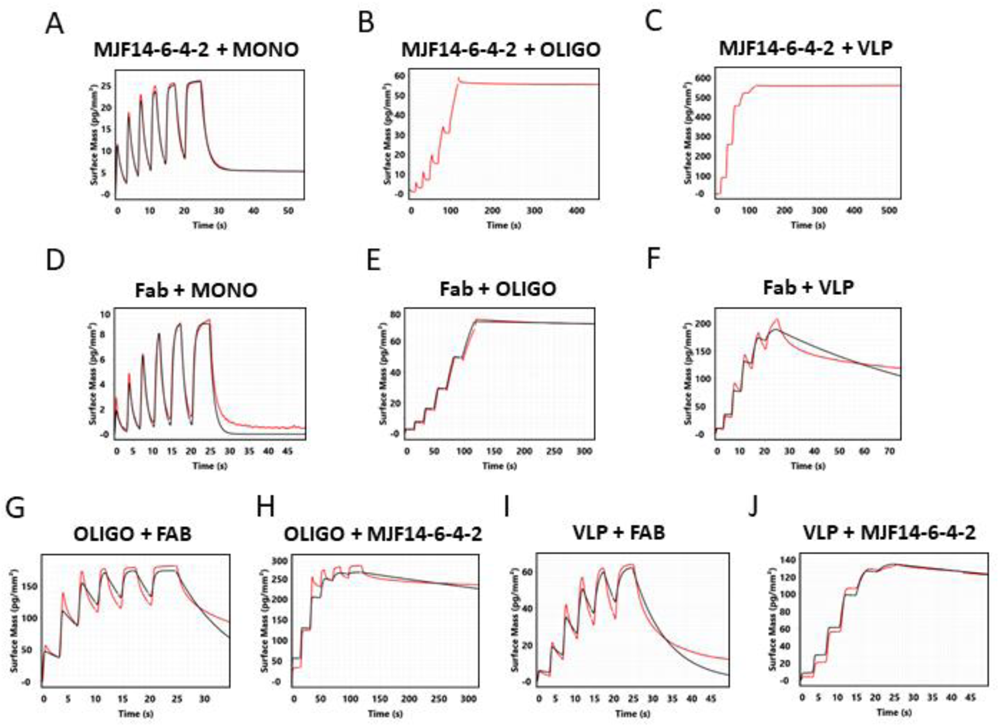
Grating-coupled interferometry (GCI) sensorgrams obtained using the WaveRAPID experiment. Sensorgrams with the MJFR14-6-4-2 antibody immobilised and **(A)** α-synuclein monomer, (**B**) α-synuclein oligomers, or (**C**) EPEA-displaying VLP applied as analytes. Sensorgrams with the MJFR14-6-4-2 Fab fragment immobilised and (**D**) α- synuclein monomer, (**E**) α-synuclein oligomers, or (**F**) EPEA-displaying VLP applied as analytes. Sensorgrams with the α-synuclein oligomers immobilised and (**G**) MJFR14-6-4-2 Fab fragment or (**H**) antibody applied as analytes. Sensorgrams with the VLP immobilised and (**I**) MJFR14-6-4-2 Fab fragment or (**J**) antibody applied as analytes. All data were double-referenced against a buffer and a reference channel. Raw, reference-subtracted data are shown in red, while fitted models are shown in black. Fitted kinetic parameters are shown in Table S2.

Control experiments with AP205 VLP showed no significant response, confirming the specificity of the interaction for the epitope (Figure S1). We then performed similar experiments with the Fab immobilised instead via amine coupling (Figure 5D-F). Here, α- synuclein monomers again showed the weakest binding (K_D_ = 2.861 μM), whereas oligomers did not dissociate from the surface. Surprisingly, with the Fab immobilised, EPEA-displaying VLP had faster dissociation and affinity in the mid-nanomolar range, contrary to observations by ELISA. This may stem from limited diffusion and binding of large species on the 3D chip surface compared to an ELISA plate. To mitigate potential surface avidity-induced effects, we performed reversed GCI measurements with α-synuclein oligomers and VLP immobilised and the antibody/Fab pair as ligands (Figure 5G-J). Here, we observed much tighter binding of the antibody to both oligomers and VLP compared to the Fab fragment. The binding of the Fab to both oligomers and VLP displayed similar mid-nanomolar binding (K_D_=184 and 136 nM, respectively), whereas the MJFR14-6-4-2 antibody had sub-nanomolar affinities. Unfortunately, in our hands, monomeric α-synuclein was not amenable to immobilisation on the GCI chip surface. Overall, GCI results indicated higher reactivity of MJFR14-6-4-2 antibodies and Fab fragments towards oligomers and VLPs than monomers.

### Crystal structure of MJFR14-6-4-2 Fab fragment in complex with epitope peptide

To further investigate the binding of antibodies, we solved the crystal structure of Fab fragments in the presence of 7-mer peptide, corresponding to C-terminal residues 134-140 of α-synuclein at 1.8 Å resolution. The electron density was clearly visible for the residues 137-140 interacting with CDR loops of Fab fragment (Figure 6). Somewhat weaker density could be observed for residues preceding Tyr 136, but no density for residues 134-135. All 4 residues from the EPEA epitope were involved in contacts with CDR loops, while Tyr 136 was not involved in any interactions apart from crystal contacts discussed further in the text. The C-terminal carboxyl group was buried in a pocket (Figure 6b), explaining why the C-terminally extended peptide EPEAG (AP-19k described in Figure 1a) containing the epitope sequence did not bind to antibodies. Five bridging water molecules were involved in epitope-Fab H-bond interactions. Somewhat surprisingly, while all 3 negative charges (two glutamates and C-terminus) took part in H-bond interactions with CDR loops, none of them was involved in salt bridge formation. This contrasts with interactions of the same epitope with nanobodies, where the C-terminal alanine was also buried, but its charge was countered by arginine from the CDR loop [32]. It should be noted that the exact recognition mode of EPEA epitope by nanobody is completely different from that of MJFR14-6-4-2.

**Figure 6.**
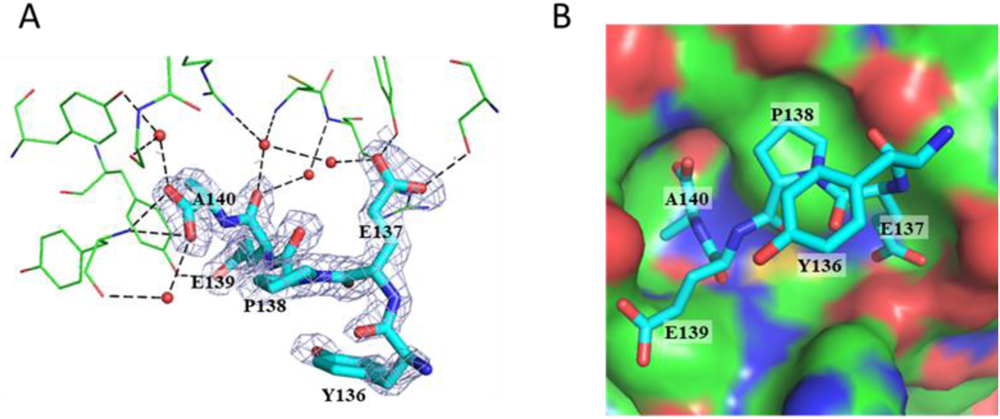
Structure of epitope peptide – Fab fragment complex. (A) Polar interactions between peptide (thick stick model) and Fab fragment residues (thin sticks). H-bonds are shown as dashed lines. Water molecules, involved in bridging interaction between Fab fragment and peptide are shown as spheres. (B) Interactions of the peptide with Fab fragment, shown as an accessible surface model. Atom colours in both panels – carbon-green (Fab) or magenta (peptide), oxygen -red, nitrogen – blue.

The crystal contacts in our structure included the interaction of the bound peptide with the neighboring Fab fragment as shown in Figure S2. This could be interpreted as a “second binding site” since each Fab fragment in the crystal has contact with two peptide molecules. In the same way, each peptide molecule is in contact with two Fab fragments. However, the “second binding site” produces only 158 Å^2^ buried surface and 2 H-bonds with Fab, while in the “main binding site”, there are 6 H-bonds and the buried surface is 341 Å^2^, almost half of the total accessible surface area of the peptide (802 Å^2^). One of two H-bonds in the “second binding site” is made by Tyr 136, which is not part of the minimal epitope. Also, while the “main binding site” is clearly located in between CDR loops, the “second binding site” merely contacts one of the CDR loops. Therefore, the “second binding site” most likely is just a crystallization artifact and does not represent additional, weaker binding site outside the crystal context.

### Solution NMR structure of epitope within monomer and VLPs

In order to compare the MJFR14-6-4-2 epitope conformation between monomer and VLP, we performed solution NMR spectroscopy on uniformly ^13^C, ^15^N-labeled samples. The ^1^H-^15^N HSQC spectrum of VLP (Figure 8A) showed 11 backbone amide correlation peaks, presumably stemming from the mobile glycine-serine linker and the epitope. Whereas the bacteriophage AP205 coat protein peaks were invisible due to the slow tumbling of the VLP resulting in peak broadening beyond solution NMR detection limits. The low 1H chemical shift dispersion suggested that the observed residues were unstructured and that a few peaks correspond well to a subset of the spectrum of monomeric, intrinsically disordered α-syn (the spectral overlay is shown in Figure S3). Using 3D NMR chemical shift assignment experiments we identified all ^15^N and ^13^C MJFR14-6-4-2 epitope backbone resonances, with the other 4 linked to the Gly-Ser linker. The back-bone walk for chemical shift assignments in 2D strips is shown in Figure S4. As anticipated, the MJFR14-6-4-2 epitope fragment with partial linker retained flexibility, evident in the 2D 1H-15N spectrum (Figure 7A). To confirm the lack of any secondary structure propensity within the epitope, we calculated the secondary chemical shifts (using the equation ΔδCα-ΔδCβ, [33]) that are sensitive indicators of protein secondary structures. Both, VLP and α-syn monomers with the same aligned residues, showed only small propensities to secondary structure elements, suggestive of random coil conformations (Figure 7B). The largest chemical shift difference deviation from zero was observed for Glu residue in both samples (Figure 7B), which was preceded by Pro residue. This is due to Pro propensity to modify Cα and Cβ chemical shifts, which are induced by conformational alterations [34]. Thus, the NMR data indicate no structure for the MJFR14-6-4-2 epitope fragment irrespective of being presented within the VLPs or on the C-terminus of monomer α-syn.

**Figure 7.**
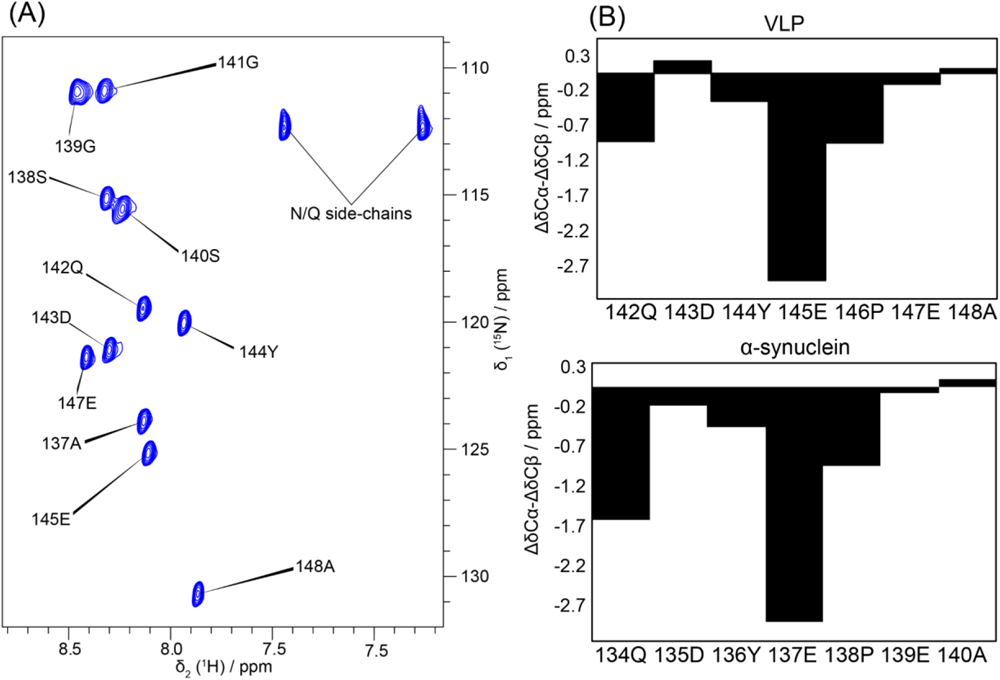
NMR data of QDYEPEA peptide within VLPs and α-synuclein. (A) The 2D ^1^H-^15^N HSQC spectrum of the VLP. Only peaks from the C-terminal part of the sample were observed. (B) The plot of secondary chemical shift vs sequence for QDYEPEA stretches of the VLP (top) and full-length α-synuclein (bottom). In these plots, the negative value on the y-axis indicates a propensity towards beta sheets, while the positive value is a propensity towards α-helices.

**Figure 8.**
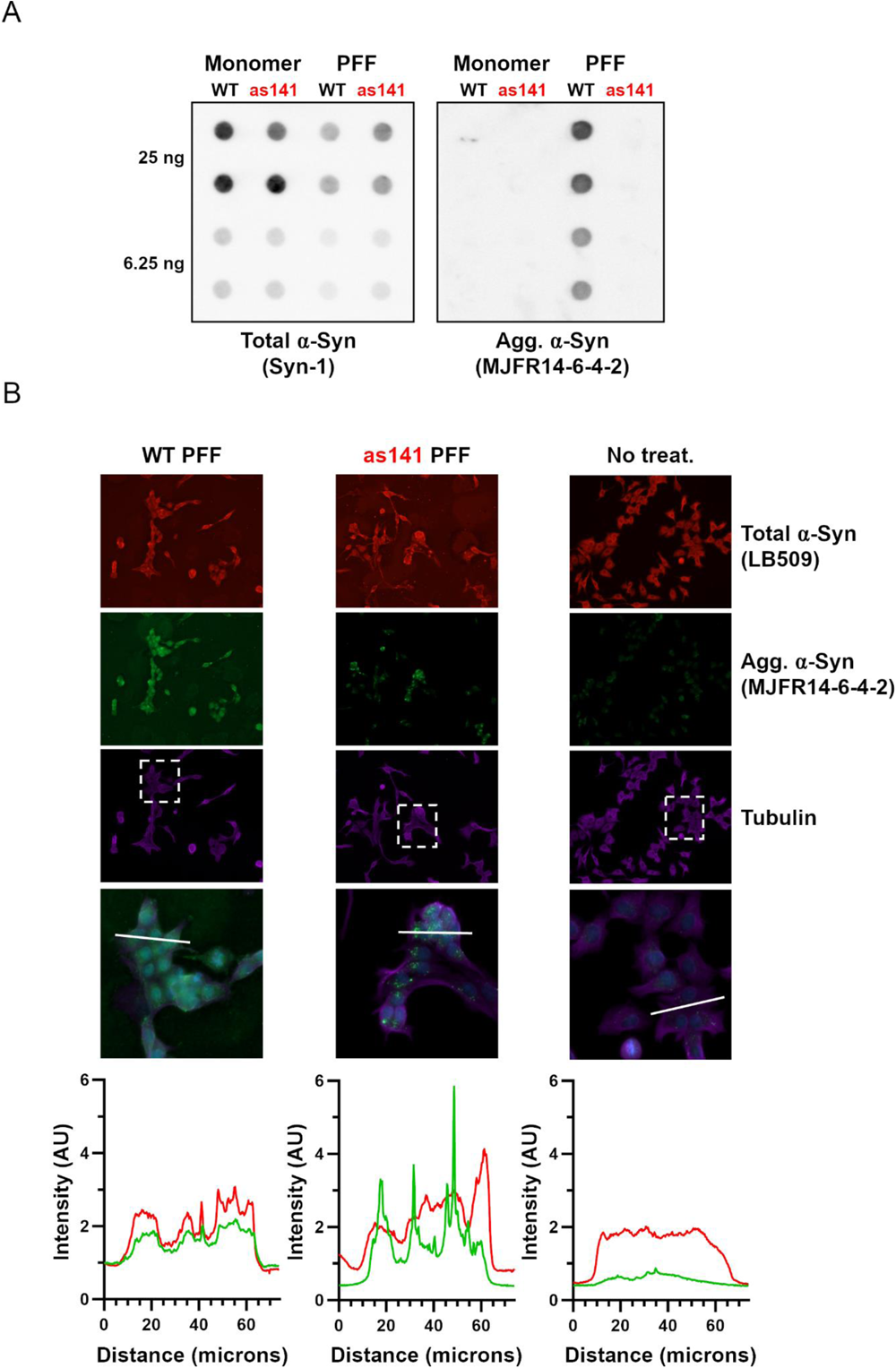
⍺-Syn with an extra C-terminal glycine is not detectable by the aggregate specific antibody MJFR14-6-4-2. PFFs were generated from human ⍺-Syn-S129A (WT) and ⍺-Syn-S129A with a C-terminal added glycine (as141). a) Native dot blotting of 25 ng, 6.25 ng WT/as141 PFF, and monomer ⍺-Syn-S129A was tested for their binding of conformation-independent antibody Syn-1 (total aSYN) and the aggregate-specific antibody MJFR-14-6-4-2. b) Rat oligodendroglial cells (OLN-AS7) stably expressing human ⍺-Syn was incubated for 24 h with WT or as141 PFF at a concentration of 14 µg/mL or PBS as negative control (No treat.) before fixation and Immunofluorescence microscopy using antibodies against tubulin (purple), aggregated ⍺-Syn (MJFR14-6-4-2, green) and ⍺-Syn (LB509, red). Note the high background signal of the Syn-1 in both PFF-treated conditions compared to the control. By contrast, the MJFR14-6-4-2 antibody only displayed a background signal in the WT-PFF treated condition whereas the as141-PFF treated sample exhibited as low a background as the non-treated control. Regions of interest in the three conditions are highlighted as hatched boxes in the tubulin image and these regions are displayed in the lower panel. Here a line scan of the pixel intensity is performed along the line displayed. The line scans measured as the grey value for the three conditions for both the Syn1 signal (red line) and MJFR14-6-4-2 signal (green line) are presented in the lowest row. Evidently, the WT-PFF treated cells display a fairly strong signal of about 1 outside the cells for both antibodies. By contrast, the as141-PFF displays a Syn1 signal as the WT-PFF treated cells, but their MJFR14-6-4-2 signal outside the is as low as the non-treated cells. Moreover, they display an enhanced signal-to-background ratio and more distinct punctae for the intracellular inclusions.

### Development of preformed α-synuclein filaments that cannot be recognized by MJFR14-6-4-2 for disease modelling

Disease modelling of α-synuclein aggregate pathophysiology in cell cultures or whole brains in vivo is at present to a large extend conducted by seeding the aggregation of intracellular α-synuclein by treating the model with exogenous preformed α-synuclein fibrils [20]. The detection of α-synuclein aggregates is often performed using antibodies toward the phospho-Ser129 epitope as a proxy because this post-translational modification correlates well with aggregates present in cellular inclusions and we have thus employed S129A mutant fibrils to avoid any detection of any phosphorylation of the preformed fibrils [20, 35]. The use of aggregate-specific antibodies like MJFR14-6–4-2 carries the risk that the exogenous preformed fibrils are recognised by the antibody thus decreasing the signal-to-noise ratio when studying the endogenously formed aggregates.

Our demonstration that adding a glycine residue to the C-terminal alanine completely blocks the MJFR14-6-4-2 binding, encouraged us to produce human recombinant α-synuclein that would be “invisible” to MJFR14-6-4-2. This should be achieved by extending the S129A expression vector with a glycine to yield human α-synuclein-S129A-141G. We successfully expressed and purified this modified construct alongside native human α-synuclein. Fibrils were generated from the wild-type (wt) α-synuclein and the S129G-141G modified α- synuclein. Using the dot blotting technique, the results reveal that the MJFR14-6-4-2 antibody only binds to preformed fibrils comprising wt-α-synuclein, while it does not bind to the S129A-141G variant fibrils, nor to either wt-or S129A-141G monomers (Figure 8A). We verified consistent loading of α-synuclein across the individual dot blots using the conformation-independent α-synuclein antibody Syn1.

To test the wt-and S129A-141G-preformed fibrils (PFF) ability to seed inclusions of α- synuclein in cells, we analyzed OLN-AS7 rat oligodendroglial cells that stably express human α-synuclein after 24 h treatment with the two variants of preformed fibrils (Figure 8b). Staining of the cells with the LB509 antibody, which binds all α-synuclein irrespective of being endogenous or the two exogenous PFF variants, demonstrates cellular staining of all conditions. Definitive inclusions are difficult to discern due to the mixed signal from both exogenous and endogenous α-synuclein. However, there is as expected no signal from the surface of the substratum used to grow the cells in the non-treated control cells in contrast to the PFF-treated conditions. The MJFR14-6-4-2 antibody intensely labelled inclusions in cells treated with both wt-and S129A-141G-preformed fibrils whereas there is no signal in the PBS controls not treated with filaments (Figure 8B). However, a green haze appears between the cells and to some from the cells that likely stems from residual PFF adsorbed to the surface of the substratum and the cells. This background staining was essentially abolished when using the S129A-141G PFF (Figure 8B). To visualise this better signal-to-noise ratio obtained by using the S129A-141G PFF in combination with the MJFR14-6-4-2 antibody, we made line scans across the inclusion-containing cells treated with wt and S129A-141G variant PFF (Figure 8B, lower row). This demonstrated the signal-to-noise ratio was higher for the S129A-141G variant indicative of less background staining (Figure 8B). By contrast, staining of the cells with the Syn-1 revealed a similar staining of cells treated with WT and S129A-141G PFF and an equal positive background staining of the substratum that contrasted to the black background of cells not treated with PFF. This demonstrates the S129A-141G PFF is superior to WT-PFF for studying seeded intracellular α-synuclein aggregation visualised by MJFR14-6-4-2 immunostaining because the exogenous S129A-141G-PFF are not detected by this particular antibody. Hence, we named the S129A-141G PFF as Stealth PFF. We recommend a similar strategy using stealth PFF be adopted in cellular modelling of seeded α-synuclein aggregation to be detected by MJFR14-6-4-2. The strategy is analogous to the use of PFF mutated on S129 because seeded cellular aggregates become phosphorylated on S129. Detection of the pS129 positive inclusions with phospho-specific antibodies will by this approach also not detect the exogenous PFF, but the pS129 staining is prone to background due to the presence of physiological nuclear pS129-α-synuclein, and the potentially late appearance of the phosphorylation in some aggregates [35, 36]. Hence, the S129A-141G Stealth PFF may be advantageous for cell biological applications but also for in vivo experiments where it is important to assure no binding to the exogenous PFF.

### Implications for drug design against synucleinopathies

Passive immunization with antibodies targeting aggregated forms of α-synuclein is a valid strategy to combat PD and other synucleinopathies, which has been evaluated in both preclinical disease models [37–41] and clinical trials [42, 43]. Although the epitope profile of these antibodies is well known, the mechanistic basis for preferential binding to aggregated α-synuclein species is not always completely understood. Our determined structure of the MJFR14-6-4-2 Fab fragment with the epitope peptide showed that the antibody bound to the C-terminal residues 137–140 and NMR studies indicated that the epitope assumed a disordered conformation. This suggests that similarly to the binding of Cinpanemab to the N-terminal epitope on α-synuclein [41], MJFR14-6-4-2 preferentially recognizes multimeric species not because of their specific conformation but due to avidity effects arising from high local concentration of epitopes, fast dissociation kinetics and low affinity towards monomers. The high avidity binding to aggregates thus emerges as a common mechanism for antibodies targeting both, N-and C-terminal epitopes of α-synuclein, which can be exploited for future drug design.

## Materials and methods

### Production of α-synuclein

Production and purification of regular and ^13^C and ^15^N labeled α-syn was performed as described previously [44, 45]. α-syn oligomers were prepared as previously described [46].

### Cloning, expression, and purification of α-synuclein-141G

As141 fibrils were generated by adding the sequence for glycine on the c-terminal of an existing human S129A ⍺-Syn construct by overhang-PCR. The amplified construct was ligated into a pET 11D vector following transformation into E. coli DH5a cells. After verifying successful cloning through sequencing, the construct was introduced into E. coli BL21(DE3) for protein expression. Protein was purified as previously described [20].

### Generation of preformed ⍺-Syn fibrils (PFF)

PFF was generated essentially as described [20]. Lyophilized wt ⍺-Syn or ⍺-Syn-141G (as141) was reconstituted in PBS pH 7.4 (Gibco) and as a novel procedure filtered through a 100 kDa filter (Amicon) to remove eventual oligomer species. Protein concentration was measured by BCA and the concentration was adjusted to 4 mg/mL. The reaction was incubated at 37 °C with continuous shaking at 1050 rpm for 72 h. The generated PFFs were harvested by centrifugation at 15,600 g for 30 min and resuspended in PBS. The protein concentration of the PFFs was adjusted and aliquots were stored at room temperature until use.

### Production and purification of phage AP205 VLP library with α-synuclein sequences

Plasmid constructs encoding bacteriophage AP205 coat protein with C-terminally attached α-synuclein peptide sequences were designed as shown in Figure 1A. Gene sequences were ordered and cloned in pETDuet-1 plasmid by BioCat GmbH, Germany. The corresponding VLPs were produced in E.coli BL21(DE3) strain and purified by gel filtration and ion exchange essentially as described before for similar VLPs [47].

### ELISA

96 well microtiter plate (#655001, Greiner, Germany) wells were coated with 1 µg in 100 µL of purified VLPs, α-syn oligomers, or monomers in technical duplicates. After blocking with 1% BSA (Sigma-Aldrich) in PBST (phosphate-buffered saline and 0,1 % Tween20), 100 µL of MJFR14-6-4-2 antibodies (#ab209538, Abcam) diluted 1:1000, 1:3000, 1:9000, 1:27000, 1:81000, 1:243000, 1:729000, or 1: 2187000 were added to each well. 100 µL of HRP-conjugated goat anti-rabbit IgG (1:10,000; #A8275-1ML, Sigma-Aldrich, USA) and 100 µL of o-Phenylenediamine dihydrochloride (OPD) solution (#P6912, Sigma-Aldrich, USA) with H2O2 were added into the wells, and the binding was detected at 492 nm absorbance on ELISA plate reader (BDSL Immunoskan MS 355, Finland).

### Competition ELISA

α-synuclein oligomers (1 μg/mL) or AP205-PD3 VLPs (1 μg/mL) were adsorbed on ELISA plates o/n at +4 °C in carbonate buffer. Wells were blocked with 1% BSA in PBST. MJFR 14-6-4-2 antibodies (20 ng/100 μL) or Fab fragments (40 ng/100 μL) were pre-incubated with 3-fold decreasing concentrations of corresponding 700 pM inhibitor (monomers, oligomers, or AP205 VLPs with epitope inserts, molar concentration calculated for amount of epitopes) in blocking solution overnight at +4 °C. 100 μL of pre-incubated MJFR 14-6-4-2 antibodies-inhibitor or Fab-inhibitor solutions were added to ELISA plate wells and incubated for 1 hour at +37 °C. 100 µL of HRP-conjugated goat anti-rabbit IgG (1:10,000; #A8275-1ML, Sigma-Aldrich, USA) and 100 µL of o-Phenylenediamine dihydrochloride (OPD) solution (#P6912, Sigma-Aldrich, USA) with H_2_O_2_ were added into the wells, and the binding was detected at 492 nm absorbance on ELISA plate reader (BDSL Immunoskan MS 355, Finland). In competition ELISA with liposome-incubated α-synuclein and VLPs, the starting concentration of inhibitors (monomers and AP-19c VLPs) was 14 nM.

### Preparation, crystallization, and structure determination of Fab fragments in complex with peptide

N-terminally acetylated peptide with sequence QDYEPEA was ordered from Metabion, Germany. Fab fragments from MJFR14-6-4-2 antibodies (#ab209538, Abcam) were prepared with Pierce™ Fab Preparation Kit (#44985, Thermo Scientific, USA) according to manufacturers’ instructions. The obtained Fab fragments were dialyzed against 20 mM Tris-HCl pH 8.0 and concentrated to 7 mg/ml by an Amicon 10 kDa spin filter cartridge. For crystallization sitting droplets were set up at 20 °C by mixing 1 μL of Fab fragments, 0.5 μL of peptide (1 mg/mL in 20 mM Tris-HCl, pH 8.0), and 1 μL crystallization buffer (24% PEG 6000, 0.1 M Na-citrate pH 5.0). Plate-shaped crystals appeared overnight, and they were flash-frozen in liquid nitrogen using mother liquor supplemented with 30% glycerol as a cryoprotectant. 1.7 Å data were collected at Diamond Light Source synchrotron (Oxfordshire, UK), beamline i03 in Unattended Data Collection mode. Data were auto-processed at beamline by autoPROC [48]. The structure was solved by molecular replacement using rabbit monoclonal antibody R53 structure (PDB code 4ZTO) as a search model in the CCP4 program Phaser [49]. The model was built in Coot [50] and refined by Refmac [51]. Data collection, refinement, and validation statistics are shown in Table S1.

### NMR spectroscopy

For NMR experiments full-length α-synuclein and VLP MJFR14-6-4-2 epitope samples were concentrated to 5 mg/mL and 14 mg/mL in a 50 mM PBS buffer at pH=7.4. All NMR spectra were acquired on a 14 T Bruker magnet equipped with an HFCN QCI cryoprobe at 25 °C. The spectra were processed using the Bruker Topspin 3.6.1 program and analysed with the CcpNmr 2.3.1 software [52]. The backbone chemical shifts for the MJFR14-6-4-2 sample were assigned using a set of 3D HNCA, HNCO, HNCACO, and HNCACB experiments. The chemical shifts for full-length α-synuclein were taken from the BMRB database; access number 25227 [53].

### Grating-coupled interferometry (GCI)

GCI experiments were performed on a Creoptix WaveDELTA instrument. All kinetics measurements were done using PBS supplemented with 0.05% Tween-20 as a running buffer. MJFR14-6-4-2 antibody was diluted to 10 µg/ml and immobilised on a 4PCP-PAG chip to a 408 pg/mm^2^ surface density. MJFR14-6-4-2 Fab fragment was immobilised on a 4PCP chip using EDC/NHS amine coupling, to a 591 pg/mm^2^ surface density. EPEA-displaying VLPs were first biotinylated using biotin-PEG4-NHS ester (Jena Bioscience, CLK-B103), after which it was immobilised on a 4PCP-STA streptavidin chip to a final density of 58 pg/mm^2^. α-synuclein oligomers were immobilised in the same manner as the VLPs to a density of 200 58 pg/mm^2^. WaveRAPID measurements were performed using either medium or high-affinity modes with double referencing against a reference channel and a running buffer run. Data were analysed using WaveControl software (Creoptix). Dissociation crops were introduced in the final analysis of the Fab fragment applied to immobilised oligomers, as there was an irreversible bulk response on the surface that precluded fitting of the final dissociation step. For the interaction between the antibody and immobilised α-synuclein oligomers, a heterogenous ligand model was used.

### Dot blotting

Non-denaturing immuno-dot blot was used to investigate PFF epitope presentation. Wt and as141 PFF and monomer (25 ng and 6.25 ng) were applied directly onto a 0.45 µM pore size nitrocellulose membrane using a vacuum filtration system (Bio-Rad BioDot Apparatus). The membrane was blocked in 5% non-fat dry milk in TBS/Tw and then incubated overnight with primary antibody Syn-1 (BD Biosciences #610,787, 1:1000) or MJFR-14-6-4-2 (Abcam #209,538, 1:450,000). Protein was visualized with a secondary HRP conjugated antibody (Dako) followed by development with ECL (GE Healthcare) and image acquisition with a Fuji Las-3000 intelligent dark box (Fujifilm).

### Cell culture and immunofluorescence microscopy

A rat oligodendroglial cell line (OLN-AS7) stably expressing ⍺-Syn was maintained at 37 °C and 5% CO_2_ in DMEM supplemented with 10% fetal calf serum, 50 U/mL penicillin, 50 µg/mL streptomycin and for OLN-AS7 cells 0.1 mg/mL Zeocin to select for ⍺-Syn expression. For experiments, cells were seeded on poly-L-lysine coated coverslips in media without Zeocin 24 h before treatment. On the day of treatment, PFF were sonicated using a Branson SFX 250 sonifier (30 ms on, 70 ms off, 6 min total on, 70% power), and immediately added to the cell media. After 24 h incubation with PFF, the cells were washed with PBS and fixed with 4% PFA. Immunostaining was performed with the primary antibodies against; tubulin (Abcam #ab6160, 1:1000), total-⍺-Syn (Abcam #ab27766, 1:500) and aggregated-⍺-Syn MJFR14-6-4-2 (Abcam #209,538, 1:25.000) essentially as described [20]. In brief, permeabilization and blocking were done with 3% BSA, 0.1% saponin in PBS followed by 1 h incubation at room temperature with primary antibodies diluted in blocking buffer. After incubation, the cells were washed three times in 0.1% saponin in PBS, then incubated 1 h with appropriate secondary antibodies (conjugated Alexa Fluor Dyes, Thermo Fischer) and DAPI at room temperature. After incubation, the cells were washed three times, and coverslips were mounted on glass slides. Images were acquired using a Zeiss Observer.Z1. Representative images were chosen and intensity profiles across a line were generated using ImageJ.

### Treatment of α-synuclein with liposomes and CD experiments

Liposomes were prepared from DMPG and DMPC phospho-lipids as described previously [54]. Liposomes were mixed with α-synuclein in PBS (5 mg/ml liposomes, 0.5 mg/ml alpha-synuclein). The liposome-α-synuclein mixture was loaded in a 0.2 ml cuvette and measured in a Jasco J-1500 CD spectrophotometer for CD experiments. As a control for α-helical structure, we used BSA (0.5 mg/ml) in PBS.

## Data availability

Atomic coordinates of Fab fragment crystal structure in complex with epitope peptide are available in PDB, accession code 8OG0.

## Author contributions

KT, PJH, IL, KJ and AE designed the study and wrote the manuscript. IL produced VLPs and performed ELISAs, KT performed crystallography, AL performed NMR experiments, LR produced α-synuclein fibrils, HG performed cell culture, microscopy and dot-blotting, TP performed GCI experiments.

## Acknowledgments

For IL and KT State Research Program project BioMedPharm VPP-EM-BIOMEDICĪNA-2022/1-0001 and JPcofuND-2 project OligoFIT supported the study. The study was supported for PHJ by JPcofuND-2 project OligoFIT, Lundbeck Foundation grants R223-2015-4222, R248-2016-2518 for Danish Research Institute of Translational Neuroscience-DANDRITE, Nordic-EMBL Partnership for Molecular Medicine, and Michael J. Fox Foundation grant MJFF-021836. TP was supported by European Regional Development fund grant 1.1.1.5/21/A/002. We thank Diamond Light Source (UK) for access to beamline i03 (proposal mx33705). We thank you for the qualified technical assistance to Shapla Bhattacharya during CD measurements and Janis Bogans during protein chromatography.

---

**Figure S1.**
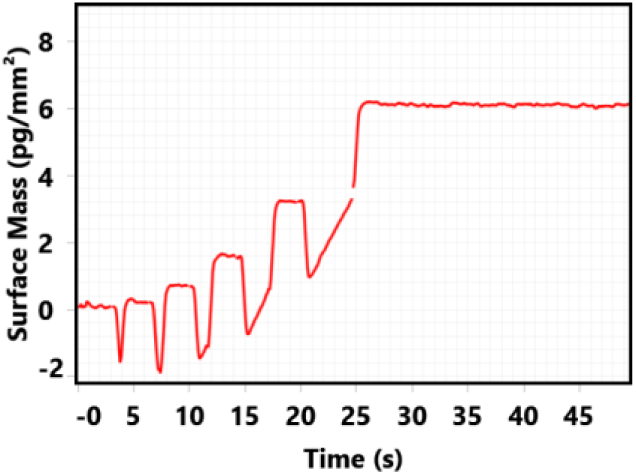
Grating-coupled interferometry (GCI) sensorgrams. obtained using the WaveRAPID experiment for the control AP205 VLP binding to immobilised MJFR14-6-4-2 antibody.

**Figure S2.**
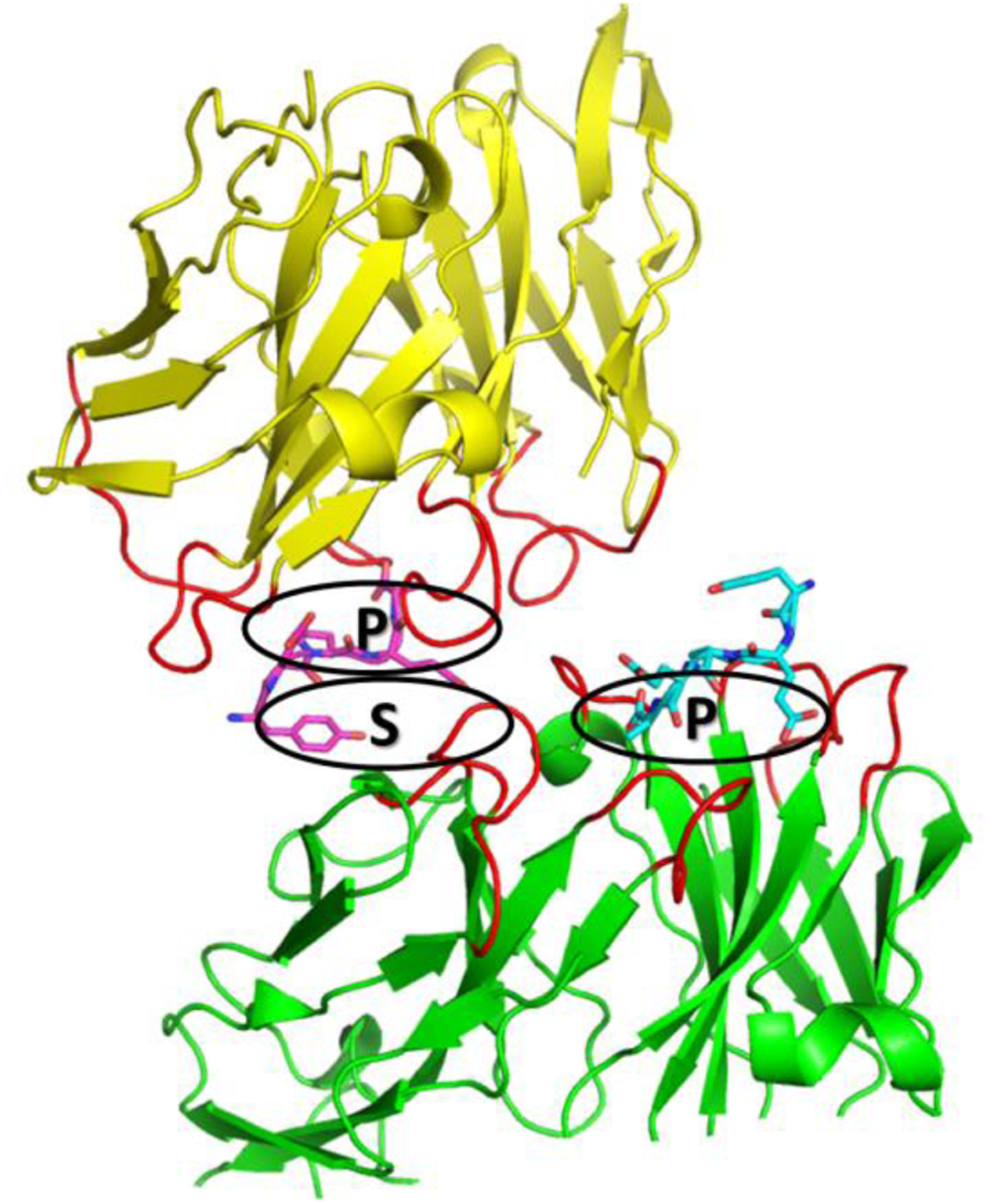
Packaging of Fab-peptide complex in a crystal lattice. The crystal contacts among asymmetric units in P2_1_2_1_2_1_ space group are such that each peptide is making interactions with two Fab fragments, variable domains of which are shown in yellow and green. The first or “primary” site, encircled as “P” is due to interactions of the peptide with a number of CDR loops (red). The second or “secondary” site, encircled as “S” is due to interactions of the same peptide with another Fab monomer in the crystal lattice.

**Figure S3.**
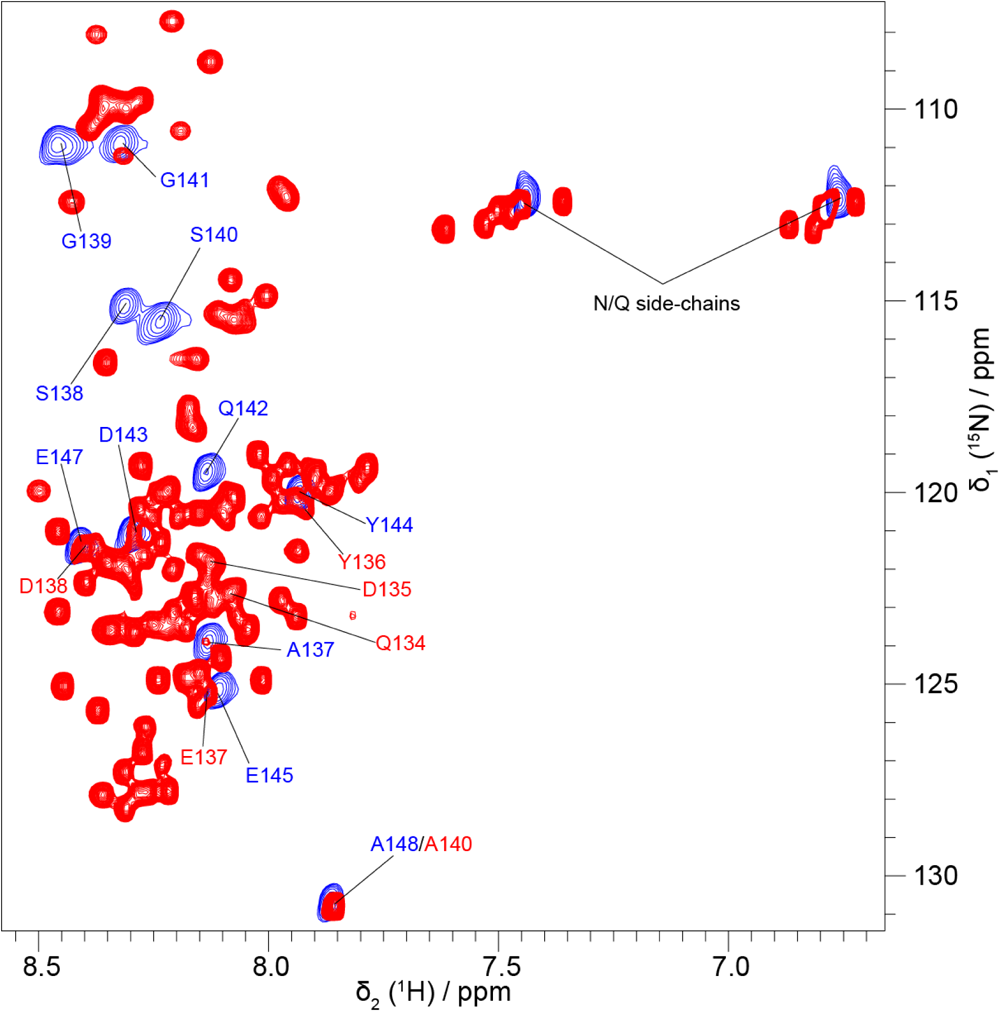
The overlay of 2D ^1^H-^15^N HSQC spectra: blue - MJFR14-6-4-2 epitope, red - full length α-synuclein. The resonance assignments only for the MJFR14-6-4-2 epitope corresponding region in α-synuclein are shown.The largest NH chemical shift deviation is for Q142 (epitope)/Q134 (α-synuclein) amino acid since its closest to the Gly-Ser linker.

**Figure S4.**
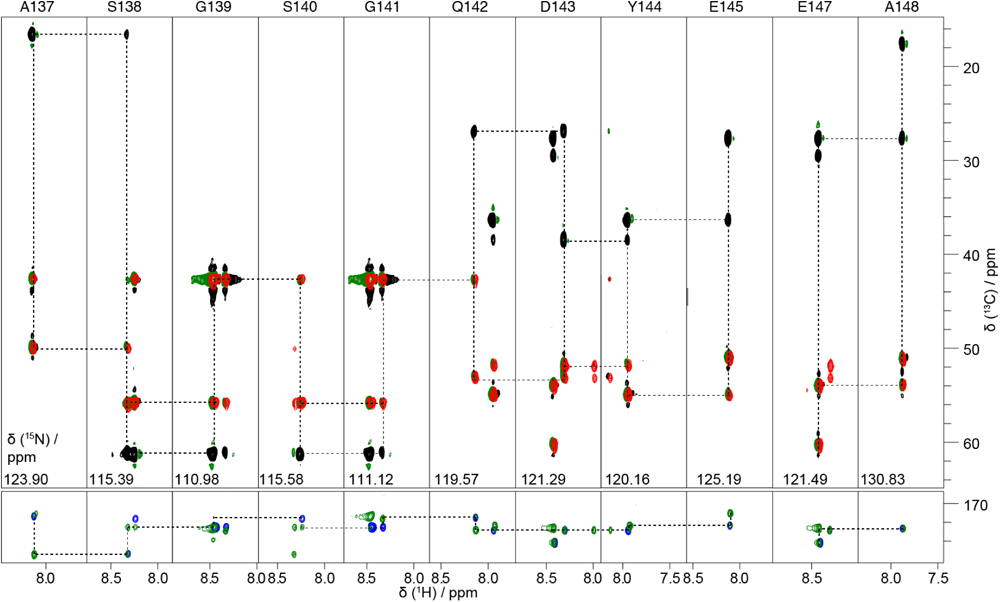
The 2D projections of 3D assignment spectra for sequence assignments of the MJFR14-6-4-2 epitope. The backbone walk for Cα_i-1_-Cα_i_, Cβ_i-1_-Cβ_i_ and CO_i-1_-CO_i_ pairs for adjacent amino acids are connected with striped lines.

**Table S1.** Data collection, refinement, and validation statistics.

| Data collection and scaling |  |
| --- | --- |
| Spacegroup | P2<sub>1</sub>2<sub>1</sub>2<sub>1</sub> |
| Cell parameters | a = 37.484 Å<br>b = 66.700 Å<br>c = 167.167 Å |
| Wavelength (Å) | 0.97625 |
| Resolution (Å) | 83.58 – 1.712 |
| Highest resolution bin (Å) | 1.902 – 1.712 |
| R<sub>merge</sub> | 0.208 (1.609) |
| Total number of observations | 425239 |
| Number of unique reflections | 32524 |
| I/σ(I) | 8.5 (1.6) |
| Completeness (%) | 87.5 (42.7) |
| Multiplicity | 13.1 (11.6) |
| Refinement |  |
| R<sub>work</sub> | 0.200 |
| R<sub>free</sub> | 0.251 |
| Average B factor (Å<sup>2</sup>) |  |
| protein | 27.6 |
| peptide | 29.6 |
| solvent | 31.3 |
| Number of atoms |  |
| protein | 3237 |
| peptide | 43 |
| solvent | 171 |
| RMS deviations from ideal |  |
| bond lengths (Å) | 0.006 |
| bond angles (°) | 1.372 |
| Ramachandran plot |  |
| residues in favored regions (%) | 96.7 |
| residues in allowed regions (%) | 99.8 |

**Table S2.** Fitted kinetic parameters for the GCI measurements.

|  | Figure 5A | Figure 5B | Figure 5C | Figure 5D | Figure 5E | Figure 5F | Figure 5G | Figure 5H | Figure 5I | Figure 5J |
| --- | --- | --- | --- | --- | --- | --- | --- | --- | --- | --- |
| Chip | 4PCP-PAG | 4PCP-PAG | 4PCP-PAG | 4PCP | 4PCP | 4PCP | 4PCP-STA | 4PCP-STA | 4PCP-STA | 4PCP-STA |
| Ligand (immobilised) | MJFR14-6-4-2 | MJFR14-6-4-2 | MJFR14-6-4-2 | Fab | Fab | Fab | OLIGO | OLIGO | VLP | VLP |
| Analyte | MONO | OLIGO | VLP | MONO | OLIGO | VLP | Fab | MJFR14-6-4-2 | Fab | MJFR14-6-4-2 |
| $k_a$, $M^{-1} s^{-1}$ | $1.2 (\pm 0.11) \times 10^6$ | No dissociation | No dissociation | $2.96 (\pm 0.13) \times 10^5$ | No dissociation | $1.00 (\pm 0.06) \times 10^5$ | $5.11 (\pm 0.49) \times 10^5$ | $2.26 (\pm 0.31) \times 10^6$ | $8.45 (\pm 0.99) \times 10^5$ | $2.84 (\pm 0.16) \times 10^5$ |
| $k_d$, $s^{-1}$ | $6.37 (\pm 0.49) \times 10^{-1}$ | | | $8.48 (\pm 0.25) \times 10^{-1}$ | | $1.89 (\pm 0.17) \times 10^{-2}$ | $9.43 (\pm 1.05) \times 10^{-2}$ | $8.21 (\pm 1.79)$ | $1.16 (\pm 0.11) \times 10^{-1}$ | $3.72 (\pm 0.97) \times 10^{-3}$ |
| $K_D$, $\mu M$ | 0.530 | | | 2.861 | | 0.189 | 0.184 | | 0.136 | 0.001 |

